# The deubiquitinase (DUB) USP13 promotes Mcl-1 stabilisation in cervical cancer

**DOI:** 10.1101/2020.07.25.220996

**Authors:** Ethan L. Morgan, Molly R. Patterson, Diego Barba-Moreno, Adam Wilson, Andrew Macdonald

## Abstract

Ubiquitination is a critical regulator of cellular homeostasis. Aberrations in the addition or removal of ubiquitin can result in the development of cancer and key components of the ubiquitination machinery serve as oncogenes or tumour suppressors. An emerging target in the development of cancer therapeutics are the deubiquitinase (DUB) enzymes that remove ubiquitin from protein substrates. Whether this class of enzyme plays a role in cervical cancer has not been fully explored. By interrogating the cervical cancer data from the TCGA consortium, we noted that the DUB USP13 is amplified in approximately 15% of cervical cancer cases. We confirmed that USP13 expression was increased in cervical cancer cell lines, cytology samples from patients with cervical disease and in cervical cancer tissue. Depletion of USP13 inhibited cervical cancer cell proliferation. Mechanistically, USP13 bound to, deubiquitinated and stabilised Mcl-1, a pivotal member of the anti-apoptotic Bcl-2 family and the reduced Mcl-1 expression contributed to the observed proliferative defect. Importantly, the expression of USP13 and Mcl-1 proteins correlated in cervical cancer tissue. Finally, we demonstrated that depletion of USP13 expression or inhibition of USP13 enzymatic activity increased the sensitivity of cervical cancer cells to the BH3 mimetic inhibitor ABT-263. Together, our data demonstrates that USP13 is a potential oncogene in cervical cancer that functions to stabilise the pro-survival protein Mcl-1, offering a potential therapeutic target for these cancers.

## Introduction

Protein ubiquitination is a critical post-translation modification that is essential for the regulation of cellular homeostasis (1). The process of ubiquitination occurs through a stepwise enzymatic cascade comprising three classes of enzymes: E1 ubiquitin-activating enzymes, E2 ubiquitin-conjugating enzymes and E3 ubiquitin ligases. The functional outcome of protein ubiquitination is dependent on the type of modification (monoubiquitin or polyubiquitin) and the linkage type within the ubiquitin chain (2). This diversity of potential ubiquitin signals is able to regulate a wide range of cellular processes. For example, K48 or K11 polyubiquitin chains are primarily responsible for promoting protein degradation, whereas chains linked by M1 or K63 polyubiquitin direct the assembly of protein complexes to regulate signal transduction (3). Consequently, the deregulation of protein ubiquitination results in the promotion of diseases including cancer. For example, defects in ubiquitin-mediated proteasomal degradation can result in either the enhanced degradation of tumour suppressor proteins, or in the stabilisation of oncogenes (4).

Ubiquitination is a highly dynamic and reversible post-translational modification, and deubiquitinase enzymes (DUBs) readily cleave ubiquitin from its protein substrates (5). To date, over 100 DUBs have been identified in the human genome that are classified into seven families on the basis of the catalytic mechanism; members of the ubiquitin C-terminal hydrolases (UCHs), ubiquitin-specific proteases (USPs), ovarian tumour proteases (OTUs), the Machado-Josephin domain superfamily (MJD), the MINDY family and the ZUFSP family function as cysteine proteases, whereas JAB1/MPN/MOV34 metalloenzymes (JAMMs) are zinc-dependent metalloproteases (5). In the last 10 years, DUBs have been shown to regulate the actions of proteins involved in cancer progression, including the tumour suppressors p53 and PTEN, and the oncogenes c-Myc and EGFR (6–9). Therefore, the pharmacological targeting of DUBs offers a potential therapeutic opportunity for treating cancers (4). For example, FT671 is a highly specific, non-covalent small molecule inhibitor of USP7, a DUB that regulates MDM2 levels, a critical negative regulator of the p53 tumour suppressor (10,11). FT671 efficiently destabilised MDM2 in Multiple Myeloid MM.1S cells, resulting in increased p53 expression and inhibiting tumour growth in murine xenografts (11).

Cervical cancer is the fourth most common cancer in women, causing significant morbidity and mortality worldwide (12). Persistent infection with human papillomavirus (HPV) is the underlying cause of almost all cervical cancers, with the high-risk HPV16 and HPV18 types being the two most prevalent HPV types responsible (13). HPV-induced transformation is primarily driven by the E6 and E7 viral oncogenes (14,15), and this is achieved through the extensive manipulation of host cell signalling networks (16–21). Critically, E6 and E7 promote the proteasomal degradation of the p53 and pRb tumour suppressors, by co-opting the E3 ligases E6-associated protein (E6-AP) and a Cullin-2 ubiquitin ligase complex (22–24). Additionally, the minor HPV oncoprotein E5 can enhance EGFR signalling (25–28), in part by disrupting the c-Cbl/EGFR complex, thereby decreasing c-Cbl-mediated ubiquitination and degradation of the EGFR (29). Several DUBs are also targeted during the HPV life cycle and in HPV-associated cancers (30–33). These data suggest that the deregulation of protein ubiquitination plays a key role in HPV associated cancers.

To identify novel DUBs involved in cervical cancer progression, we interrogated the cervical cancer data from the TCGA consortium. We identified that USP13 is amplified in about 15% of cervical cancer cases. We then showed that USP13 was required for the proliferation of cervical cancer cells, at least in part via the deubiquitination and stabilisation of the pro-survival protein, Mcl-1. Crucially, we demonstrated that pharmacological inhibition of USP13 sensitises HPV+ cervical cancer cells to BH3 mimetic inhibitors, suggesting that targeting of USP13 may have therapeutic benefit in these cancers.

## Results

### Increased USP13 expression in pre-malignant cervical disease and cervical cancer

To investigate whether DUBs contribute to transformation in cervical cancer, we analysed the cervical cancer dataset from The Cancer Genome Atlas (TCGA) (34). By examining the copy number alterations (CNAs) of all the known DUB genes in the human genome, we identified amplification of *USP13* in approximately 15 % of cervical cancers, as well as in a number of other squamous carcinomas (Fig 1A). Importantly, *USP13* copy number positively correlated with *USP13* mRNA expression (R=0.350) (Fig. 1B). Next, we examined the expression of USP13 in a panel of cervical cancer cell lines. When compared to primary normal human keratinocytes (NHKs), *USP13* mRNA expression was higher in HPV positive (HPV+), but not HPV negative (HPV-) cervical cancer cells (Fig. 1C). USP13 protein levels were also increased in HPV+ cervical cancer cells compared with HPV-cervical cancer cells, or NHKs, when analysed by western blot (Fig 1D). To confirm the increased USP13 protein expression in cervical cancer, we performed immunohistochemistry (IHC) on a cervical cancer tissue microarray (TMA). In line with the data from cell lines, USP13 protein expression was significantly higher in the cervical cancer tissue (Fig. 1E).

**Figure 1.**
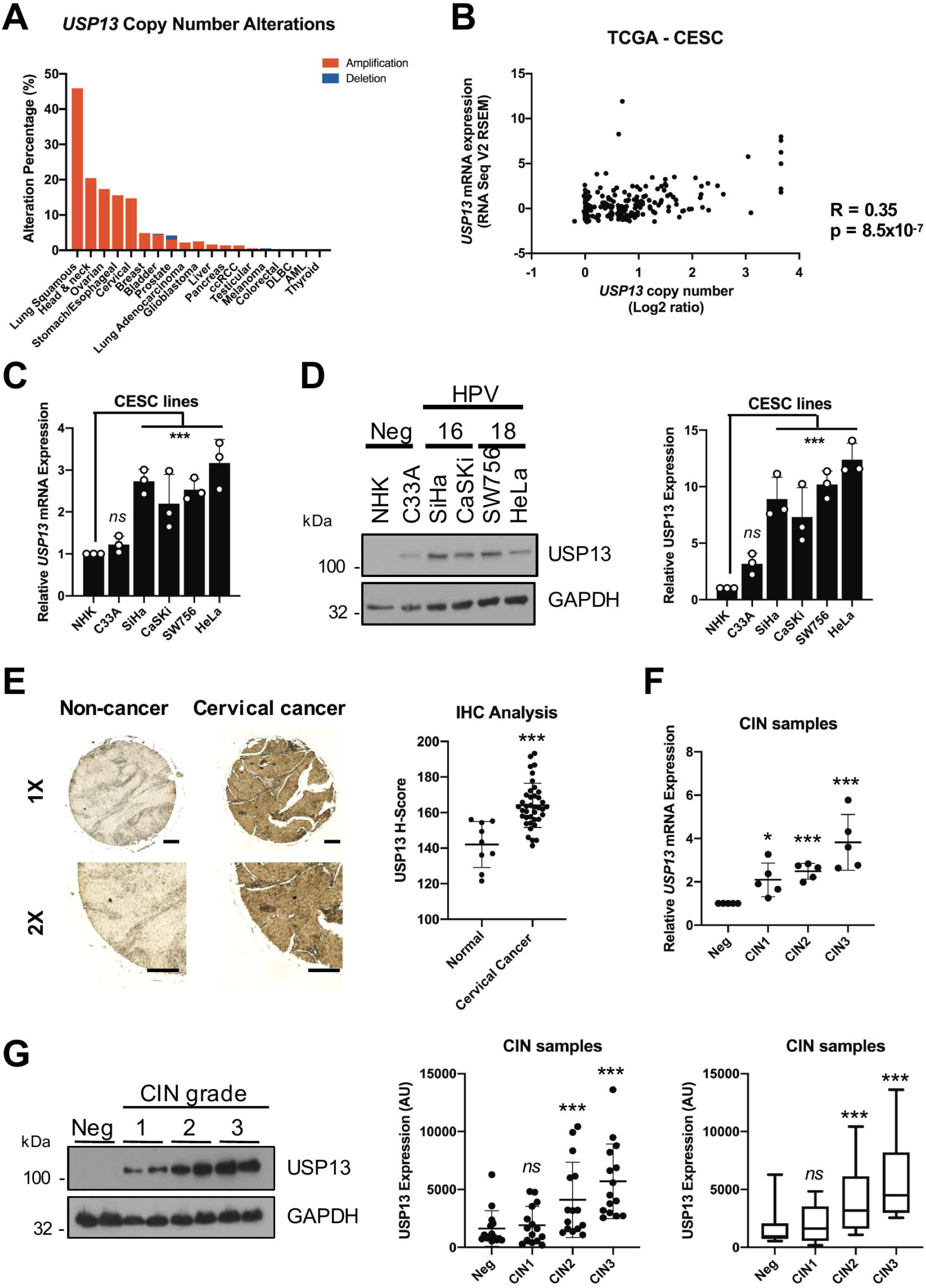
USP13 expression is upregulated in pre-malignant cervical disease and cervical cancer. **A)** Genomic alterations of *USP13* across human cancers determined by cBioportal analysis of TCGA data. **B)** Scatter dot plot analysis of *USP13* mRNA expression against *USP13* copy number alterations in cervical cancer determined by cBioportal analysis of TCGA data. Correlation was determined using Spearman’s analysis. **C)** RT-qPCR analysis of *USP13* mRNA expression in normal human keratinocytes (NHKs), HPV-C33A cells, HPV16+ SiHa and CaSKi cells and HPV18+ SW756 and HeLa cells. mRNA expression was normalized against *U6* mRNA levels. **D)** Representative western blot of USP13 expression in normal human keratinocytes (NHKs), HPV-C33A cells, HPV16+ SiHa and CaSKi cells and HPV18+ SW756 and HeLa cells. GAPDH served as a loading control. Quantification of the protein band intensities from four biological, independent repeats are shown on the right. **E)** Representative immunohistochemical (IHC) staining of USP13 expression in cervical cancer tissues and normal cervical epithelium from a tissue microarray (TMA). Scale bars, 10025FBμm. Scatter dot plot analysis of USP13 expression from a larger cohort of cervical cancer cases (n=41) and normal cervical epithelium (n=9) is shown on the right. **F)** Scatter dot plot of RT-qPCR analysis of *USP13* mRNA expression from a panel of cervical cytology samples representing CIN lesions of increasing grade. Five samples from each clinical grade (negative (Neg) and CIN I-III) were analysed and mRNA levels were normalized to the negative samples. Samples were normalized against *U6* mRNA levels. **G)** Representative western blot of cervical cytology samples of CIN lesions of increasing grade analysed for USP13 protein expression. GAPDH served as a loading control. Scatter dot and box plot analysis of a larger cohort of samples (n=15 for each grade) is shown on the right. Bars represent the mean ± standard deviation from at least three biological repeats unless otherwise stated. * p<0.5; ** p<0.01; *** p<0.001 (Student’s t-test).

The development of cervical cancer occurs over many years, through the accumulation of pre-malignant alteration of the squamous epithelia collectively known as cervical intraepithelial neoplasia (CIN); CIN1 represents a transient HPV infection with mild dysplasia, while CIN3 represents severe dysplasia which may develop into cervical cancer (35). To investigate if USP13 expression may contribute to the development of cervical cancer, we analysed *USP13* mRNA expression in cervical cytology samples from a cohort of HPV16+ patients. Samples from healthy, HPV-patients were used as controls. *USP13* mRNA expression and the levels of USP13 protein both increased during progression through CIN1 to CIN3 (Fig.1F and 1G). In validation of our data in cervical cancer cell lines and cervical cancer tissue, *USP13* mRNA expression was also significantly upregulated in several published microarray databases (Supplementary Fig. 1), suggesting that increased USP13 expression is a common occurrence in cervical cancer.

### USP13 expression is regulated by c-Jun/AP-1 in cervical cancer cells

In addition to the *USP13* amplification observed by our analysis of the TCGA cervical cancer dataset (Fig. 1A), we noted that a number of cases had high *USP13* mRNA expression in the absence of *USP13* amplification. We therefore wished to further investigate how *USP13* expression is regulated in cervical cancer. Whilst oncogenes can be upregulated in cancer at the transcriptional level, the transcriptional regulation of USP13 is poorly understood. To gain insight, we analysed the putative *USP13* promoter region upstream of the *USP13* start codon for transcription factor binding sites (Fig. 2A). We identified two AP-1 like sequences that have previously been shown to bind to c-Jun, a member of the AP-1 transcription factor family that is highly active in cervical cancer (36,37). To test if c-Jun/AP-1 regulates *USP13* expression we first inhibited the activity of the MAP kinase (MAPK) JNK, which is an important inducer of c-Jun/AP-1 signalling (38). Addition of the potent, specific irreversible JNK inhibitor JNK-IN-8 abrogated c-Jun phosphorylation and reduced c-Jun expression, as previously observed. Furthermore, we saw a substantial reduction in USP13 protein expression (Fig. 2B). JNK inhibition also decreased *USP13* mRNA expression, demonstrating that JNK-mediated c-Jun activity may regulate USP13 at the transcriptional level (Fig. 2C). To investigate if c-Jun directly played a role, we depleted c-Jun expression using a pool of specific siRNAs. Knockdown of c-Jun resulted in a significant decrease in *USP13* mRNA and protein expression in both HeLa and SiHa cells (Fig. 2D and E). Finally, we assessed if c-Jun regulated USP13 expression directly, by binding the *USP13* promoter via the putative AP-1 binding sites. We performed chromatin immunoprecipitation (ChIP) analysis utilising primers covering the AP-1 like sequences in the *USP13* promoter (Termed AP-1-1 and AP-1-2; Fig. 2A). In DMSO treated HeLa cells, there was a significant enrichment of c-Jun binding to the AP-1-1 site in the *USP13* promoter compared to an IgG isotype control. In contrast, little to no binding of c-Jun was observed at AP-1-2 (Fig. 2F). Crucially, c-Jun binding to the AP-1-1 site was dependent on JNK activity, as treatment with JNK-IN-8 abolished c-Jun binding to the *USP13* promoter. Taken together, JNK-dependent c-Jun activation is necessary for USP13 expression in cervical cancer cells.

**Figure 2.**
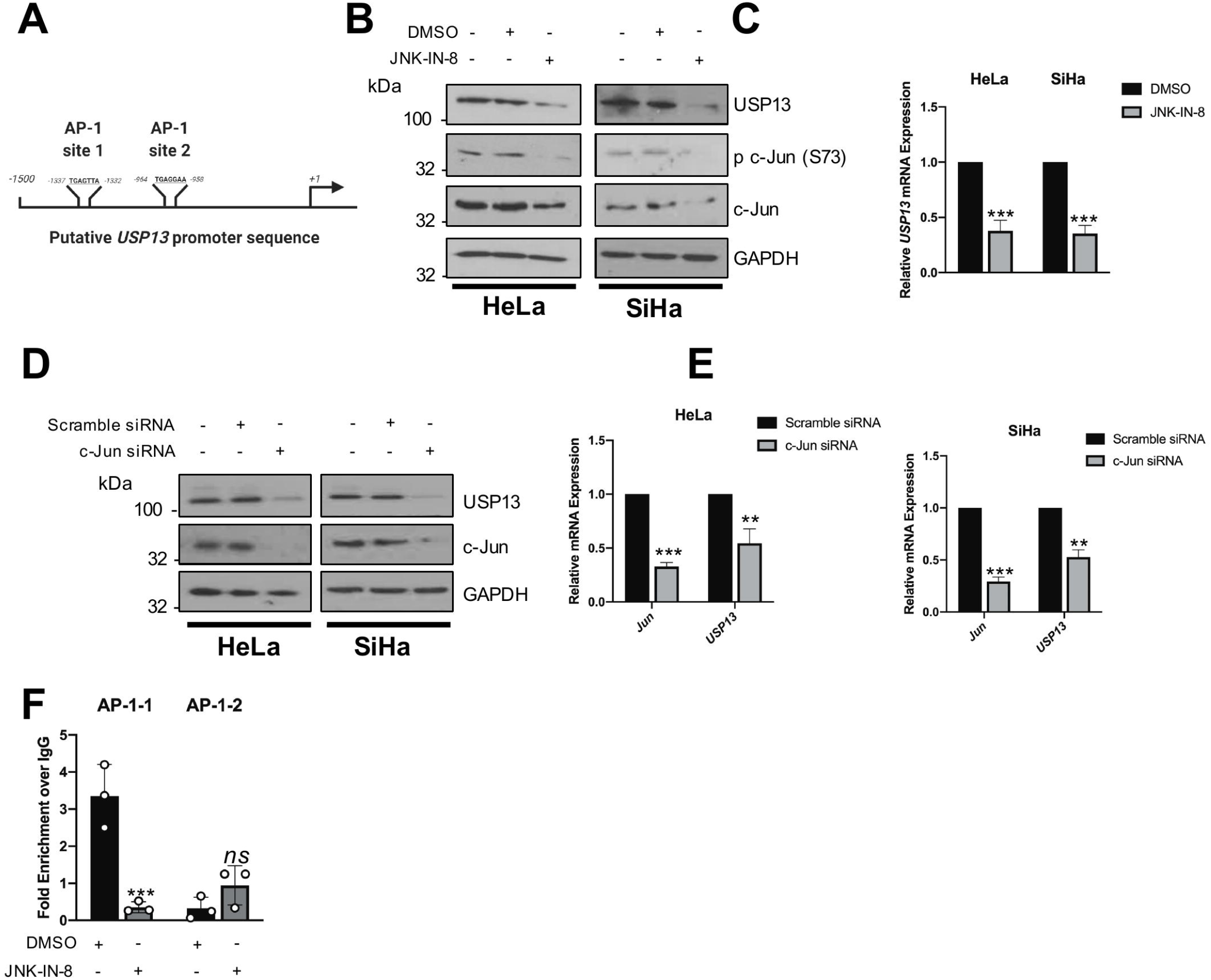
USP13 expression is regulated by c-Jun/AP-1 activity in cervical cancer cells. **A)** Schematic of potential AP-1 binding sites in the *USP13* promoter region. **B)** Representative western blot of HeLa and SiHa cells after treatment with JNK-IN-8 (10 μM) or DMSO control for 48 hours. Lysates were analysed for the expression of USP13, phosphorylated c-Jun (S73) and c-Jun. GAPDH was used as a loading control. **C)** RT-qPCR analysis of *USP13* mRNA expression in HeLa and SiHa cells after treatment with JNK-IN-8 (10 μM) or DMSO control for 48 hours. mRNA expression was normalized against *U6* mRNA levels. **D)** Representative western blot of HeLa and SiHa cells after transfection of a pool of four specific c-Jun siRNA for 72 hours. Lysates were analysed for the expression of USP13 and c-Jun. GAPDH was used as a loading control. **E)** RT-qPCR analysis of *USP13* and c*JUN* mRNA expression of HeLa and SiHa after transfection of a pool of four specific c-Jun siRNA for 72 hours. mRNA expression was normalized against *U6* mRNA levels. **F)** Chromatin was prepared from HeLa cells and c-Jun was immunoprecipitated using an anti-c-Jun antibody, followed by RT-qPCR using primers specific to the two putative AP-1 binding sites in the *USP13* promoter. c-Jun binding is presented as percentage of input chromatin. Bars are the means ± standard deviation from at least three biological repeats. * p<0.5; ** p<0.01; *** p<0.001 (Student’s t-test).

### USP13 is required for cervical cancer cell proliferation

Depletion of USP13 using a pool of specific siRNAs (Fig. 3A) caused a significant reduction in cervical cancer cell growth (Fig. 3B). The absence of USP13 also led to a reduction in colony formation under anchorage dependent (Fig. 3C) and independent conditions (Fig. 3D). Conversely, over-expression of wild type (WT) USP13 enhanced cell growth and colony formation (Fig. 3E-H). This enhancement was dependent on USP13 deubiquitinase activity, as a catalytically inactive mutant of USP13 (USP13 C345A) failed to increase cell growth. Together, these results show that expression of catalytically active USP13 is required for the proliferation of cervical cancer cells.

**Figure 3.**
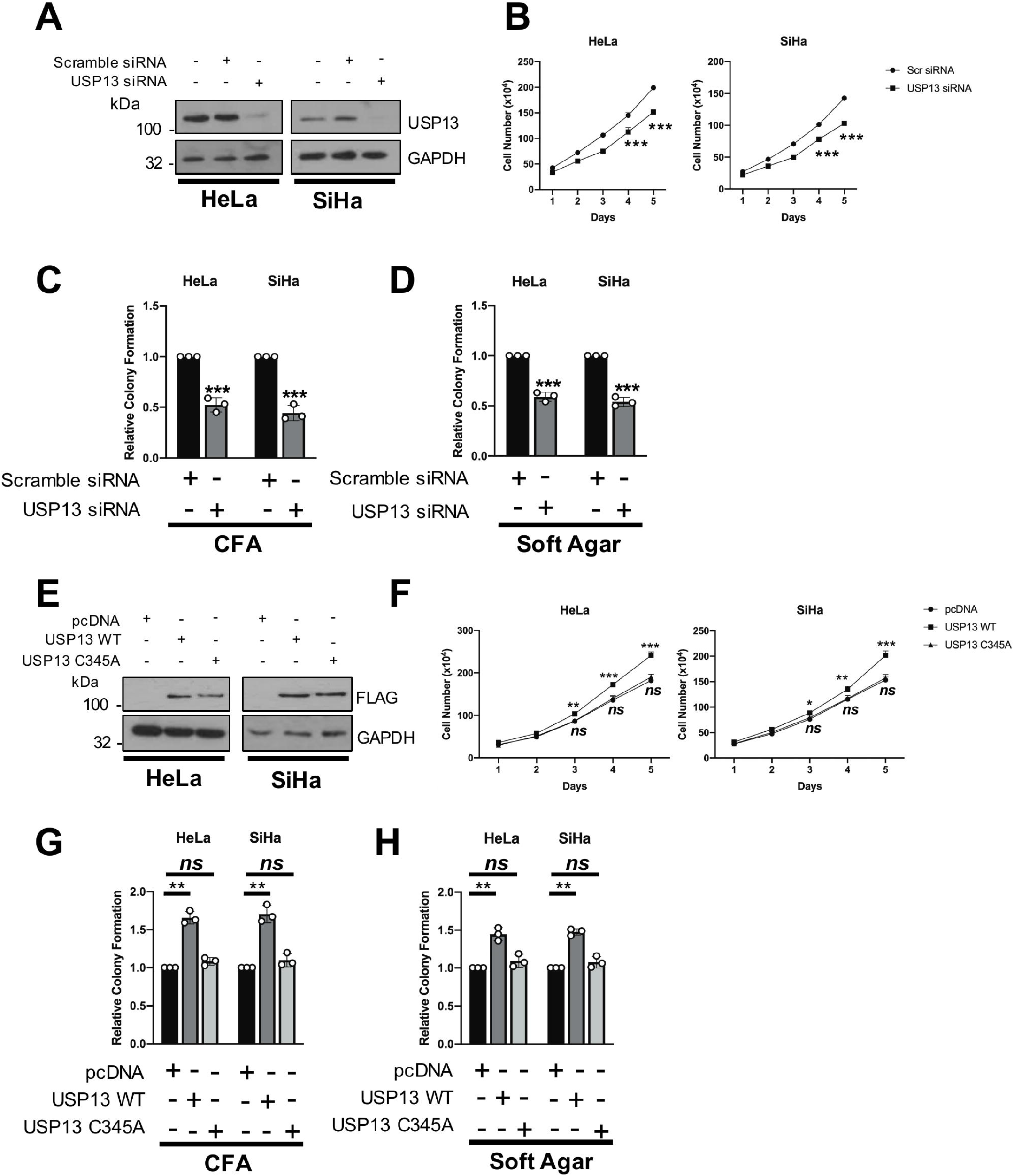
USP13 expression is required for the proliferation of cervical cancer cells. **A)** Representative western blot of HeLa and SiHa cells after transfection of a pool of four specific USP13 siRNA for 72 hours. Lysates were analysed for the expression of USP13 and GAPDH was used as a loading control. **B)** Growth curve analysis of HeLa and SiHa cells after transfection of a pool of four specific USP13 siRNA for 72 hours. **C)** Colony formation assay (anchorage dependent growth) of HeLa and SiHa cells after transfection of a pool of four specific USP13 siRNA for 72 hours. **D)** Soft agar assay of HeLa and SiHa cells after transfection of a pool of four specific USP13 siRNA for 72 hours. **E)** Representative western blot of HeLa and SiHa cells after transfection of a FLAG-USP13 or FLAG-USP13 (C345A) for 48 hours. Lysates were analysed for the expression of USP13 and GAPDH was used as a loading control. **F)** Growth curve analysis of HeLa and SiHa cells after transfection of a FLAG-USP13 or FLAG-USP13 (C345A) for 48 hours. **G)** Colony formation assay (anchorage dependent growth) of HeLa and SiHa cells after transfection of a FLAG-USP13 or FLAG-USP13 (C345A) for 48 hours. **H)** Soft agar assay of HeLa and SiHa cells after transfection of a FLAG-USP13 or FLAG-USP13 (C345A) for 48 hours. Bars are the means ± standard deviation from at least three biological repeats. * p<0.5; ** p<0.01; *** p<0.001 (Student’s t-test).

### USP13 promotes Mcl-1 stability

Several USP13 targets have been identified, including microphthalmia-associated transcription factor (MITF), ATP citrate lyase (ACLY) and phosphatase and tensin homolog (PTEN) (39–41). Interestingly, USP13 was recently shown to deubiquitinate and stabilise the pro-survival protein Mcl-1 in lung and ovarian cancers (42). Like cervical cancer, ovarian cancers are squamous cell carcinomas; therefore, we hypothesised that USP13 might also regulate Mcl-1 in cervical cancer cells. To investigate this, the level of Mcl-1 expression was analysed in USP13 depleted cells. Loss of USP13 significantly reduced Mcl-1 protein expression, without effecting *MCL1* mRNA expression (Fig. 4A and B). Critically, treatment of USP13 depleted cells with the proteasome inhibitor MG132 rescued Mcl-1 protein levels, suggesting that USP13 protects Mcl-1 from proteasomal degradation (Fig. 4C). As a complementary approach, the over-expression of USP13 caused an increase in Mcl-1 protein levels (Fig. 4D). In agreement with our proliferation data, over-expressed catalytically inactive USP13 did not increase Mcl-1 protein levels (Fig. 4D). Again, USP13 over-expression had no effect on *MCL1* mRNA expression (Fig. 4E), suggesting that the regulation of Mcl-1 by USP13 is post-transcriptional.

**Figure 4.**
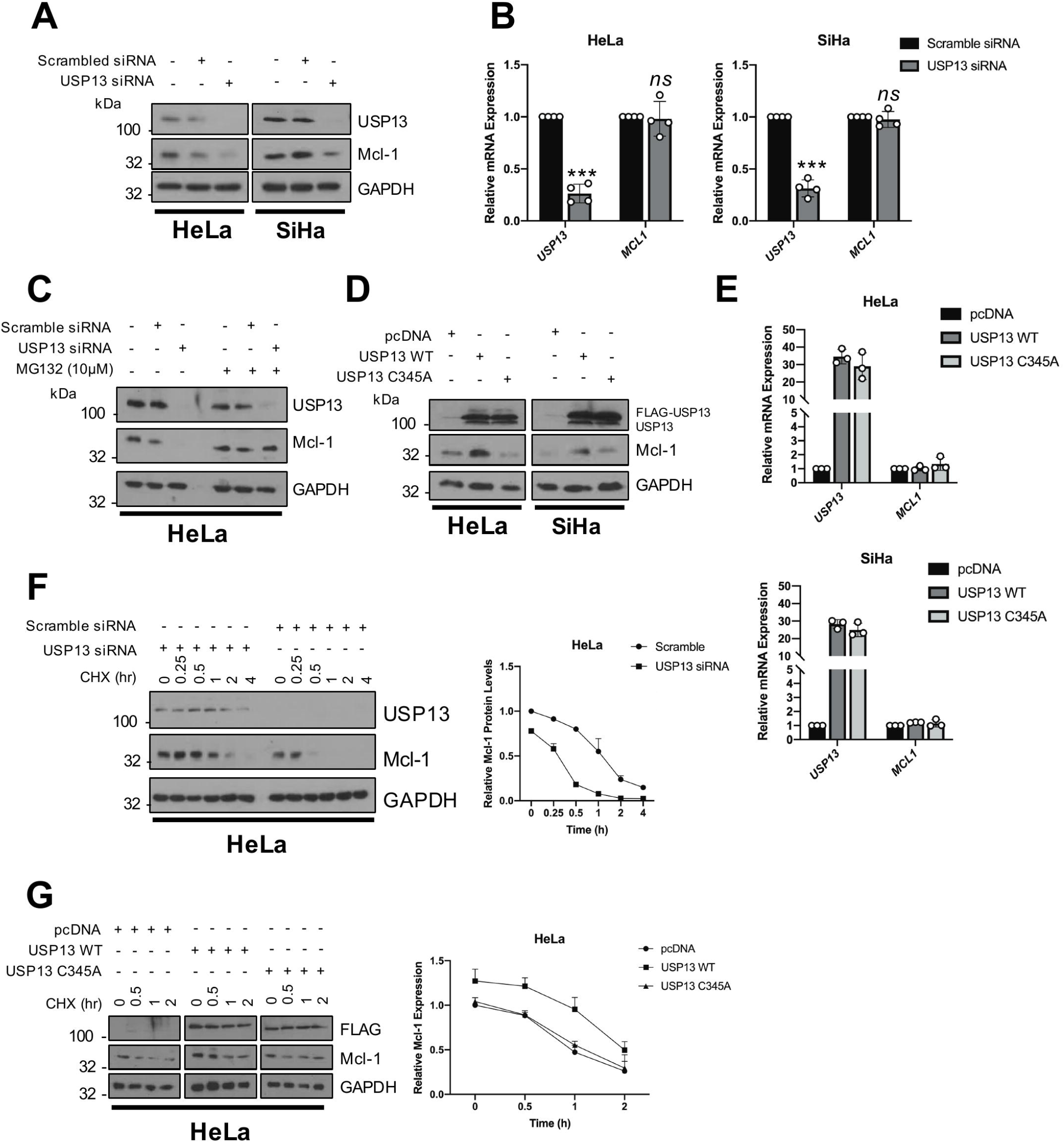
USP13 promotes the stability of the pro-survival protein Mcl-1. **A)** Representative western blot of HeLa and SiHa cells after transfection of a pool of four specific USP13 siRNA for 72 hours. Lysates were analysed for the expression of USP13 and Mcl-1. GAPDH was used as a loading control. **B)** RT-qPCR analysis of *USP13* and *MCL1* mRNA expression of HeLa and SiHa cells after transfection of a pool of four specific USP13 siRNA for 72 hours. mRNA expression was normalized against *U6* mRNA levels. **C)** Representative western blot of HeLa and SiHa cells after transfection of a pool of four specific USP13 siRNA for 72 hours. Cells were additionally treated with 10 μM MG132 for 6 hours. Lysates were analysed for the expression of USP13 and Mcl-1. GAPDH was used as a loading control. **D)** Representative western blot of HeLa and SiHa cells after transfection of FLAG-USP13 or FLAG-USP13 (C345A). Lysates were analysed for the expression of USP13 and Mcl-1. GAPDH was used as a loading control. **E)** RT-qPCR analysis of *USP13* and *MCL1* mRNA expression of HeLa and SiHa cells after transfection of FLAG-USP13 or FLAG-USP13 (C345A). mRNA expression was normalized against *U6* mRNA levels. **F)** Representative western blot of HeLa cells after transfection of a pool of four specific USP13 siRNA for 72 hours. Cells were additionally treated with 20 μM cycloheximide and harvested at the indicated time points. Lysates were analysed for the expression of USP13 and Mcl-1. GAPDH was used as a loading control. Quantification of the protein band intensities from three biological, independent repeats are shown on the right. **G)** Representative western blot of HeLa cells after transfection of FLAG-USP13 or FLAG-USP13 (C345A). Cells were additionally treated with 20 μM cycloheximide and harvested at the indicated time points. Lysates were analysed for the expression of USP13 and Mcl-1. GAPDH was used as a loading control. Quantification of the protein band intensities from three biological, independent repeats are shown on the right. Bars are the means ± standard deviation from at least three biological repeats. * p<0.5; ** p<0.01; *** p<0.001 (Student’s t-test).

The effect of USP13 on Mcl-1 protein turnover was investigated by cycloheximide chase assay. When USP13 was depleted from HeLa cells, the half-life of Mcl-1 was markedly reduced, from approximately 90 mins to approximately 20 mins (Fig. 4F). Conversely, overexpression of WT USP13, but not the USP13 mutant, enhanced the half-life of Mcl-1 (Fig. 4G). These data together demonstrated that USP13 promotes the stability of Mcl-1 at a post-transcriptional level, by protecting it from proteasomal degradation.

### USP13 interacts with and deubiquitinates Mcl-1 in cervical cancer cells

USP13 has previously been shown to directly bind to Mcl-1 *in vitro* (42). Furthermore, Mcl-1 has also been shown to be highly expressed in cervical cancer and correlates with proliferation and survival (43). We first performed co-immunoprecipitation assays to confirm the interaction between USP13 and Mcl-1 by transfecting HEK293T cells with FLAG-USP13 and V5-hMcl-1. Immunoprecipitation with anti-FLAG antibody demonstrated that FLAG-USP13 interacted with over-expressed Mcl-1 (Fig. 5A). Furthermore, reciprocal immunoprecipitations using anti-V5 antibody confirmed this interaction (Fig. 5A). To identify if this interaction also occurred between endogenous USP13 and Mcl-1 in cervical cancer cells, immunoprecipitations were performed with either anti-USP13 or anti-Mcl-1 antibodies in HeLa and SiHa cells lysates. Immunoprecipitation with either antibody resulted in the detection of both USP13 and Mcl-1, suggesting that the endogenous proteins interact in cervical cancer cells (Fig. 5B).

**Figure 5.**
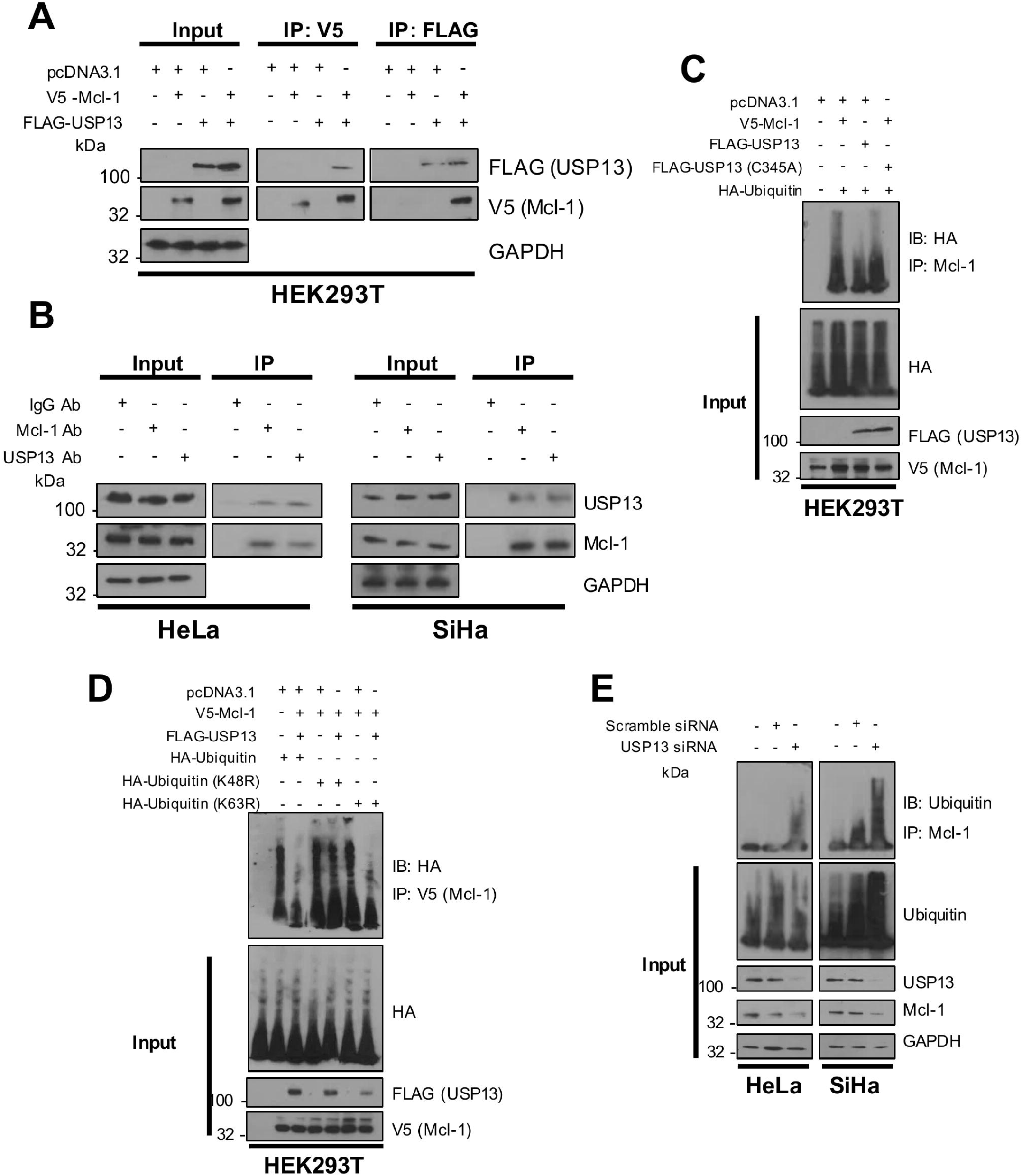
USP13 interacts with and deubiquitinates Mcl-1 in cervical cancer cells. **A)** HEK293T cells were transfected with pcDNA3.1-V5-Mcl-1, Flag-USP13, or both Mcl-1 and FLAG-USP13. Cells were treated with 10 μM MG132 for 6 hours and either Mcl-1 or USP13 were immunoprecipitated using an anti-V5 or anti-FLAG antibody. Co-immunoprecipitated Mcl-1 or FLAG-USP13 were detected using the respective antibodies. GAPDH was used as a loading control. **B)** Endogenous Mcl-1 and USP13 were immunoprecipitated from HeLa and SiHa cells after treatment with 10 μM MG132 for using an anti-Mcl1 or anti-USP13 antibodies. Co-immunoprecipitated Mcl-1 or USP13 were detected using the respective antibodies. GAPDH was used as a loading control. **C)** HEK293T cells were co-transfected with pcDNA3.1-V5-Mcl-1 and HA-Ubiquitin, with or without Flag-USP13 or a FLAG-USP13 mutant (C345A). Cells were treated with 10 μM MG132 for 6 hours and V5-Mcl-1 was immunoprecipitated using an anti-Mcl1 antibody. Ubiquitinated Mcl-1 was detected using an anti-HA antibody. GAPDH was used as a loading control. **D)** HEK293T cells were co-transfected with pcDNA3.1-V5-Mcl-1, HA-Ubiquitin or mutant Ubiquitin (K48R or K63R), with or without Flag-USP13. Cells were treated with 10 μM MG132 for 6 hours and V5-Mcl-1 was immunoprecipitated using an anti-V5 antibody. Ubiquitinated Mcl-1 was detected using an anti-HA antibody. GAPDH was used as a loading control. **E)** HeLa and SiHa cells were transfected with a pool of four specific USP13 siRNA for 72 hours. Cells were treated with 10 μM MG132 for 6 hours before harvesting. Mcl-1 was immunoprecipitated using an anti-Mcl1 antibody. Ubiquitinated Mcl-1 was detected using an anti-ubiquitin antibody. GAPDH was used as a loading control.

Next, we confirmed that USP13 deubiquitinated Mcl-1. HEK293T cells were transfected with FLAG-USP13, V5-hMcl-1 and HA-Ubiquitin, and anti-V5 immunoprecipitates were probed for the level of ubiquitination using the HA-antibody. Co-expression of USP13 and Mcl-1 led to a marked reduction in Mcl-1 ubiquitination (Fig. 5C, compare lanes 2 and 3). Consistent with our previous data, the catalytically inactive USP13 mutant was not able to reduce Mcl-1 ubiquitination, despite retaining the ability to bind to Mcl-1 (Fig. 5C, compare lanes 2 and 4; Supplementary Fig. 2). Whilst *in vitro* ubiquitination assays confirmed that USP13 is capable of directly deubiquitinating Mcl-1 (42), the precise ubiquitin linkage that USP13 removes from Mcl-1 has not been determined. Ubiquitin contains seven lysine residues that serve as sites of ubiquitination. The most studied types are K48 polyubiquitin, which targets proteins for degradation, and K63 polyubiquitin, which is predominantly associated with signal transduction (3). To determine which ubiquitin linkage USP13 deubiquitinates, HEK293T cells were transfected with the HA-tagged K48R or K63R ubiquitin mutants, which are unable to be polyubiquitinated with K48 or K63 linkages, respectively. Overexpressed USP13 was able to cleave wild-type and the K63R form of ubiquitin from Mcl-1 but not K48R ubiquitin (Fig. 5D). These results suggest that USP13 removes K48 polyubiquitin, but not K63 polyubiquitin, chains from Mcl-1. Finally, we confirmed that depletion of USP13 from both HeLa and SiHa cells increased the ubiquitination of endogenous Mcl-1, suggesting USP13-mediated deubiquitination of Mcl-1 is relevant in cervical cancer cells (Fig 5E).

### Restoration of Mcl-1 expression partially rescues the proliferation defect in USP13 depleted cervical cancer cells

As well as having a critical role in cell survival, Mcl-1 can also play an important role in cell proliferation (44). As USP13 depletion resulted in the inhibition of cell proliferation in cervical cancer cells, we investigated if this was due to the reduction in Mcl-1 levels. We therefore over-expressed Mcl-1 in USP13-depleted cells (Fig. 6A). Re-introduction of Mcl-1 resulted in a partial, but significant, restoration of cell growth (Fig. 6B) and colony formation (Fig. 6C). These results suggest that the effect of USP13 on cell proliferation in cervical cancer cells is at least partially dependent on the stabilisation of Mcl-1 expression.

**Figure 6.**
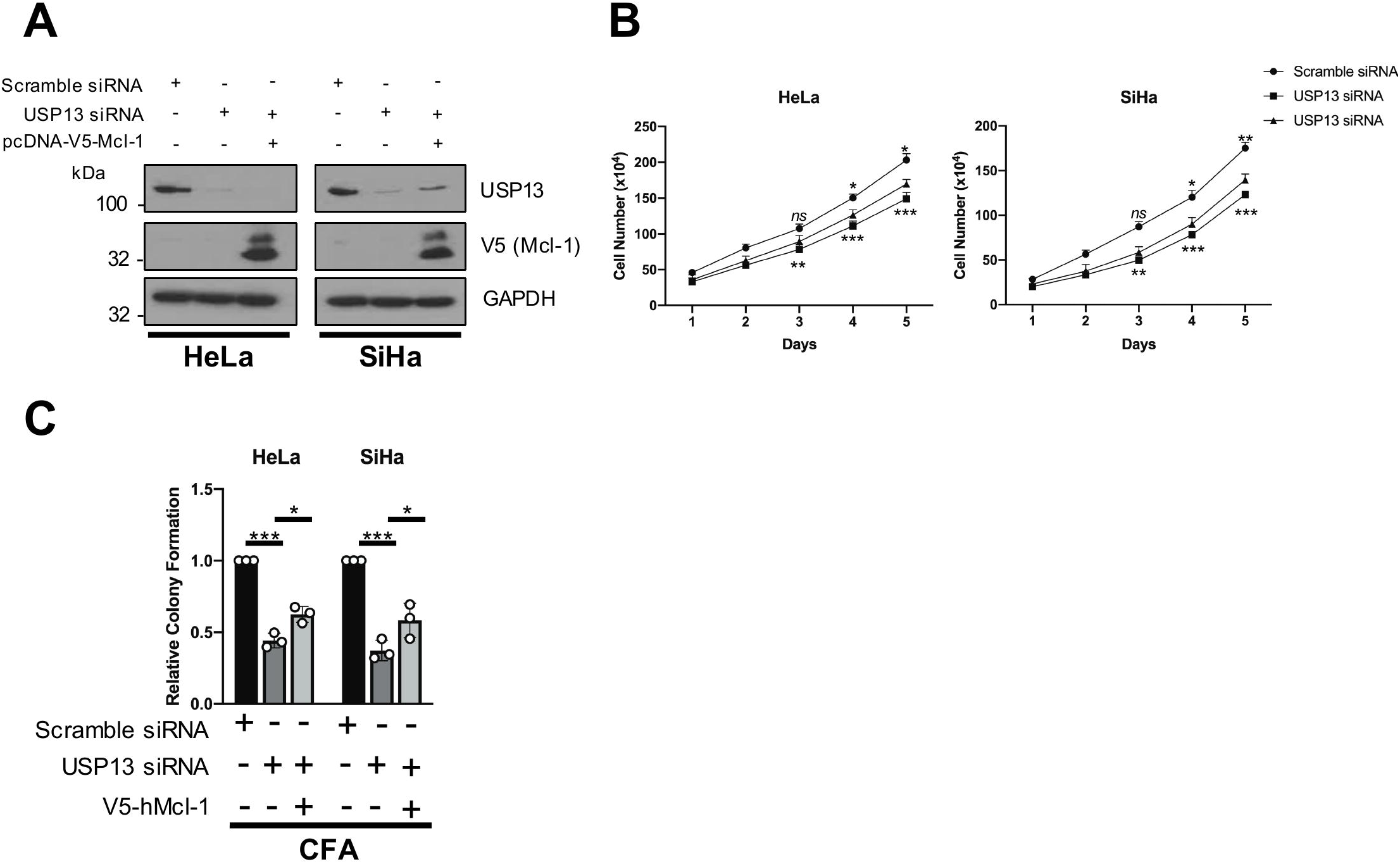
Restoration of Mcl-1 expression partially rescues the proliferation defect in USP13 depleted cervical cancer cells. **A)** Representative western blot of HeLa and SiHa cells after transfection of a pool of four specific USP13 siRNA for 72 hours. After 24 hours, cells were transfected with pcDNA3.1-hMcl-1 or control vector. Lysates were analysed for the expression of USP13, Mcl-1 and GAPDH was used as a loading control. **B)** Growth curve analysis of HeLa and SiHa cells after transfection of a pool of four specific USP13 siRNA for 72 hours. After 24 hours, cells were transfected with pcDNA3.1-hMcl-1 (triangle) or control vector (square). **C)** Colony formation assay (anchorage dependent growth) of HeLa and SiHa cells after transfection of a pool of four specific USP13 siRNA for 72 hours. After 24 hours, cells were transfected with pcDNA3.1-hMcl-1 or control vector. Bars are the means ± standard deviation from at least three biological repeats. * p<0.5; ** p<0.01; *** p<0.001 (Student’s t-test).

### USP13 and Mcl-1 expression correlate in cervical cancer

To confirm the clinical relevance of the USP13 – Mcl-1 interaction, we first analysed the expression of Mcl-1 in our panel of cervical cancer cells. Mcl-1 protein levels were significantly higher in cervical cancer cells when compared to NHKs (Fig. 7A) and correlated with increased USP13 expression in the cervical cancer cell lines (R=0.890, p=0.002). We assessed Mcl-1 expression in cervical cancer tissue by immunohistochemistry and observed higher expression of Mcl-1 in cervical cancer tissue when compared to non-cancer tissue (Fig. 7B-C). Importantly, expression of Mcl-1 significantly correlated with USP13 expression in cervical cancer tissue from the same patient (Fig 7D; R=0.4128, p=0.009). Collectively, these findings demonstrate that USP13 and Mcl-1 are concomitantly over expressed in cervical cancer tissue, suggesting that the regulation of Mcl-1 by USP13 is clinically relevant.

**Figure 7.**
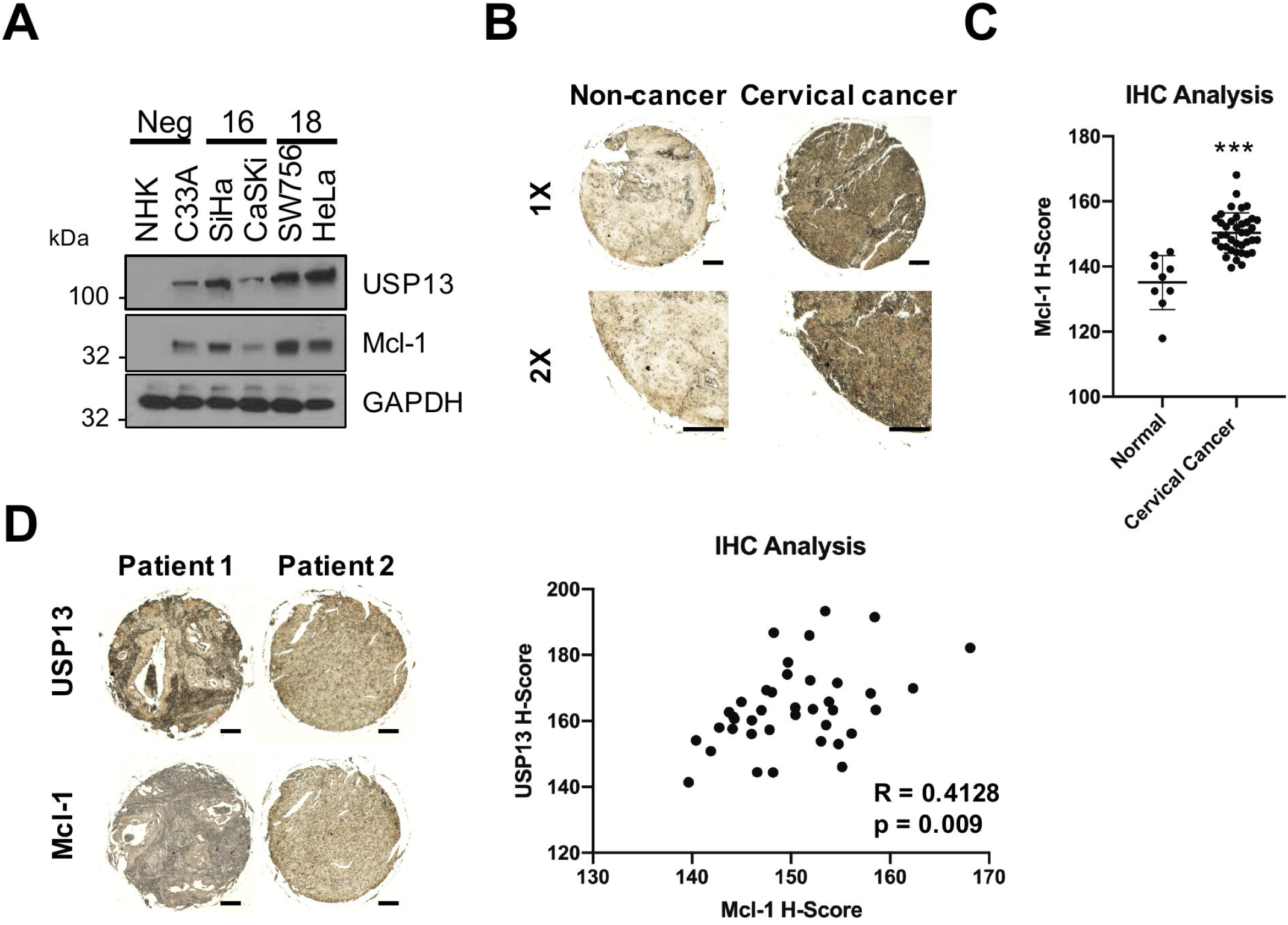
USP13 and Mcl-1 expression correlate in cervical cancer. **A)** Representative western blot of Mcl-1 expression in normal human keratinocytes (NHKs), HPV-C33A cells, HPV16+ SiHa and CaSKi cells and HPV18+ SW756 and HeLa cells. GAPDH served as a loading control. Quantification of the protein band intensities are shown on the right. **B)** Representative immunohistochemical (IHC) staining of Mcl-1 expression in cervical cancer tissues and normal cervical epithelium from a tissue microarray (TMA). Scale bars, 10025FBμm. **C)** Scatter dot plot analysis of Mcl-1 expression from a larger cohort of cervical cancer cases (n=41) and normal cervical epithelium (n=9) is shown on the right. **D)** Representative immunohistochemical (IHC) staining of USP13 and Mcl-1 expression in cervical cancer tissue from two patients. Staining was performed from separate cores from the same patient samples. Scale bars, 10025FBμm. Correlation was determined using Spearman’s analysis and is shown on the right. * p<0.5; ** p<0.01; *** p<0.001 (Student’s t-test).

### Inhibition of USP13 sensitises cervical cancer cells to BH3 mimetics

Bcl-2 family members are highly expressed in cervical cancer and inhibition of these proteins using BH3 mimetics is a promising therapeutic strategy (45). Despite this promise, over-expression of Mcl-1 is commonly associated with resistance to these inhibitors and inhibition or depletion of Mcl-1 can restore sensitivity to BH3 mimetics (46). To investigate if targeting USP13 to reduce Mcl-1 expression has therapeutic potential in cervical cancer, we first assessed the impact of USP13 inhibition by the small molecule inhibitor Spautin-1 (47). In both HeLa and SiHa cells, inhibition of USP13 by Spautin-1 resulted in a dose-dependent decrease in Mcl-1 protein expression (Fig. 8A). Spautin-1 also reduced USP13 expression, as has been previously observed in other cell lines (Fig. 8A). Similar to USP13 depletion, treatment with Spautin-1 had no effect on *MCL1* mRNA expression (Fig. 8B). Furthermore, the reduction in Mcl-1 protein expression upon Spautin-1 treatment was lessened by inhibition of the proteasome (Fig. 8C). Treatment with Spautin-1 also abolished the increase in Mcl-1 expression observed upon USP13 overexpression, suggesting that the function of Spautin-1 may be through the inhibition of USP13 (Fig 8D).

**Figure 8.**
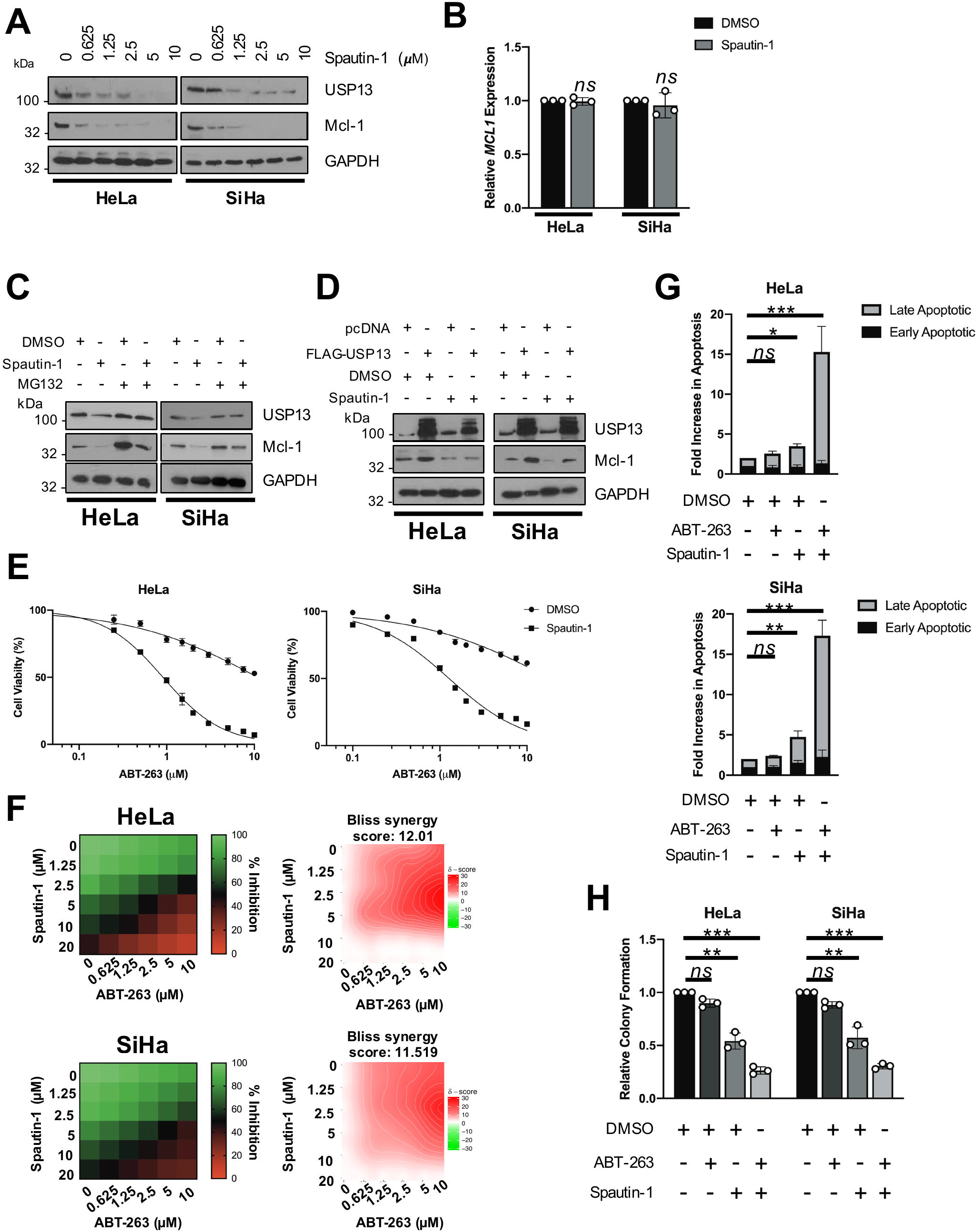
Pharmacological inhibition of USP13 sensitises HPV+ cervical cancer cells to BH3 mimetics. **A)** Representative western blot of HeLa and SiHa cells after treatment with increasing doses of Spautin-1 or DMSO control for 48 hours. Lysates were analysed for the expression of USP13 and Mcl-1. GAPDH was used as a loading control. **B)** RT-qPCR analysis of *MCL1* mRNA expression in HeLa and SiHa cells after treatment with Spautin-1 (10 μM) or DMSO control for 48 hours. mRNA expression was normalized against *U6* mRNA levels. **C)** HeLa and SiHa cells were treated with Spautin-1 (10 μM) or DMSO control for 48 hours. Cells were additionally treated with 10 μM MG132 for 6 hours. Lysates were analysed for the expression of USP13 and Mcl-1. GAPDH was used as a loading control. **D)** Representative western blot of HeLa and SiHa cells after transfection of FLAG-USP13 with or without Spautin-1 (10 μM) or DMSO control. Lysates were analysed for the expression of USP13 and Mcl-1. GAPDH was used as a loading control. **E)** Cell viability assay (MTT) of HeLa and SiHa cells treated with increasing doses of ABT-263, with or without Spautin-1 (10 μM), for 24 hours. **F)** Flow cytometric analysis of Annexin V assay in HeLa and SiHa cells after treatment with DMSO, Spautin-1 (10 μM), ABT-263 (5 μM) or both for 24 hours. **G)** Colony formation assay of HeLa and SiHa cells after treatment with DMSO, Spautin-1 (10 μM), ABT-263 (5 μM) or both for 24 hours. Error bars represent the mean ± standard deviation of a minimum of three biological repeats. * *p* < 0.05, ** *p* < 0.01, *** *p* < 0.001 (Student’s *t*-test).

Cervical cancer cells are resistant to the BH3 mimetic ABT-263 (48). We therefore investigated if pharmacological inhibition of USP13 using Spautin-1 sensitised cervical cancer cells to ABT-263. In line with previous reports, HeLa and SiHa cells were relatively insensitive to ABT-263, with an IC_50_ 10.64 and 18.58 μM, respectively ((48); Fig. 8E). However, combination treatment with Spautin-1 significantly sensitised cervical cancer cells to ABT-263 (HeLa, 12-fold increase; SiHa, 15-fold increase). To expand on this, we performed dose matrix experiments containing a range of ABT-263 and Spautin-1 concentrations and assessed potential drug synergy using the Bliss independence model (49). An overall negative Bliss synergy score indicates antagonism; a value of zero indicates additive activity, and positive Bliss synergy score indicates synergy. The heatmaps for Bliss synergy scores showed that ABT-263 in combination with Spautin-1 synergistically inhibited cell viability across multiple concentrations (Fig. 8F).

As a single agent, ABT-263 did not induce apoptosis in cervical cancer cells, whereas Spautin-1 alone did, particularly in SiHa cells (Fig. 8G). Combination treatment resulted in around a 15-fold increase in apoptosis in both cell lines (Fig. 8G). Finally, we demonstrated that whilst Spautin-1 reduced colony formation by around 50% (similar to USP13 depletion), combination treatment with ABT-263 resulted in a 75% reduction in colony formation (Fig. 8H). Taken together, these data suggest that USP13 mediated stabilisation of Mcl-1 may be responsible for the resistance of cervical cancer cells to BH3 mimetics.

## Discussion

Deregulation of the ubiquitin system is a common occurrence in diverse cancers (50). As such, targeting the enzymes responsible for either the addition or removal of ubiquitin offers a potential therapeutic avenue. Encouragingly, a number of small molecule inhibitors of ubiquitin-associated enzymes are currently entering pre-clinical trials (51–55). Here, we screened the TCGA cervical cancer database for the expression of DUBs, enzymes which catalyse the remove of ubiquitin from protein substrates (5). We identified that *USP13* is the most amplified DUB gene in cervical cancer. Interestingly, *USP13* sits on chromosome 3q26.33, in the so-called 3q26 ‘OncCassette’, a common amplicon that is found in most squamous cell carcinomas and includes the oncogene *PIK3CA* (56). Several substrates for USP13 have recently been identified; however, due to this, the function of USP13 in tumourigenesis is controversial, as it has been shown to regulate the expression of both tumour suppressors and oncogenes. USP13 can induce Beclin-1 mediated autophagy as part of a p53 regulatory loop (7) and stabilise the tumour suppressor PTEN in breast cancer (40). In contrast, USP13 also deubiquitinates and stabilizes microphthalmia-associated transcription factor (MITF), an important lineage-specific master regulator in melanoma (39). Furthermore, USP13 can deubiquitinate and stabilise ACLY/OGDH and c-Myc in ovarian cancer and glioblastoma, respectively, thereby functioning as an oncogene (41,57). Indeed, USP13 depletion did induce a proliferation defect in some ovarian cancer cells but not others, and this was dependent on the USP13 expression levels (41,42).

As well as being regulated via copy number alterations, genes can also be regulated at a transcriptional level. The cervical cancer TCGA dataset demonstrates that 8% of cervical cancers have high *USP13* mRNA expression in the absence of *USP13* copy number amplification. We therefore investigated how *USP13* expression was regulated in cervical cancer cells. We identified that the *USP13* promoter contains two AP-1-like sequences that have previously been shown to bind to the AP-1 family member c-Jun (36,37). In this study, we demonstrated that inhibition of JNK, or depletion of c-Jun, resulted in a substantial loss of USP13 expression and this was regulated at the transcriptional level (Fig. 2B-E). Furthermore, chromatin immunoprecipitation assays demonstrated that c-Jun directly bound one of the identified AP-1-like sequences in the *USP13* promoter (Fig. 2A and F). Thus, we have identified an additional mechanism of USP13 regulation.

Here, we provide evidence that USP13 functions as an oncogene in cervical cancer by promoting cell proliferation. We demonstrate that this is linked with the deubiquitination and stabilisation of the pro-survival Bcl-2 family member Mcl-1. We further show that USP13 specifically removes K48 polyubiquitin chains from Mcl-1 and these polyubiquitin chains are associated with proteasomal degradation. In lung and ovarian cancers, USP13 does not contribute to the cell proliferation of unstressed cells, whereas we observed a more general proliferative defect upon USP13 depletion (Fig. 3). Interestingly, restoration of Mcl-1 expression only partially restored cell proliferation in USP13 depleted cervical cancer cells. This suggests that additional USP13 substrates contribute to cervical cancer cell proliferation. Further studies will be required to identify these targets and understand how they contribute to USP13-mediated cell proliferation.

Over recent years, Mcl-1 has attracted attention as a potential therapeutic target in cancer (58,59). Mcl-1 plays a critical role as a pro-survival factor, as well as a pro-proliferative factor, in many tumour types. Furthermore, Mcl-1 turnover can be controlled by a number of E3 ligases (such as SCF^β-TrCP^, SCF^FBW7^ and Mule) and DUBs (such as USP9X and DUB3) (reviewed in (60)). Previously, the DUBs OTUD1 and JOSD1 had been shown to regulate Mcl-1 expression in cervical cancer cells (61,62). In contrast to the data presented here, one of these studies did not find a role for USP13 in the regulation of Mcl-1 in HeLa cells (61); however, that study focussed on overexpression of USP13. In contrast, we have taken a more holistic approach and assessed the role of USP13 in Mcl-1 stabilisation with a range of complementary assays, providing robust evidence that USP13 can indeed regulate Mcl-1 expression in cervical cancer cells.

As Bcl-2 family members have been reported to be highly expressed in cervical cancer, BH3 mimetics may have therapeutic potential in these cancers and the high expression of Mcl-1 has been implicated as a resistance mechanism to these inhibitors (45,46). Despite their promise, an inherent problem with the most advanced BH3 mimetics, ABT-236 and ABT-737, which target Bcl-2 and Bcl-_X_L, is the development of intrinsic resistance due to high expression of Mcl-1, which these inhibitors do not target (63,64). Therefore, reducing Mcl-1 expression, by direct or indirect inhibition, can sensitise resistant cancer cells to BH3 mimetics (64,66). As such, Mcl-1 specific inhibitors, such as AMG 176, can synergise with BH3 mimetics to achieve tumour regression in acute myeloid leukemia (AML) tumour models (63). Furthermore, studies with cyclin-dependent kinase (CDKs) inhibitors such as alvocidib, or multi-kinase inhibitors such as Sorafenib have shown similar results (67,68). Our data shows that USP13 and Mcl-1 protein expression correlates, suggesting that USP13-mediated Mcl-1 stabilisation is clinically relevant. Furthermore, we demonstrated that this mechanism could be exploited pharmacologically, by use of a recently identified small molecule inhibitor of USP13 to reduce Mcl-1 protein expression (7). This may have potential benefit in cervical cancer patients, as although standard chemotherapeutic options in cervical cancer have good efficacy, many cases develop local recurrences and metastases (69). Our data demonstrate that inhibition of USP13 with Spautin-1 reduces Mcl-1 expression and sensitises cervical cancer cells to the BH3 mimetic ABT-263, offering a potential therapeutic strategy for cervical cancer.

In conclusion, we have identified that the DUB USP13 is a potential oncogene in cervical cancer that promotes proliferation by deubiquitinating and stabilizing the pro-survival protein Mcl-1. We also demonstrate that pharmacological inhibition of USP13 sensitises cervical cancer cells to BH3 mimetic inhibitor by reducing Mcl-1 protein expression, inducing cell death. Therefore, we suggest that targeting USP13 may offer a therapeutic benefit in cervical cancer patients via the reduction in Mcl-1 protein expression.

## Materials and methods

### Cervical cytology samples

Cervical cytology samples were obtained from the Scottish HPV Archive (http://www.shine/mvm.ed.ac.uk/archive.shtml), a biobank of over 20,000 samples designed to facilitate HPV research. The East of Scotland Research Ethics Service has given generic approval to the Scottish HPV Archive as a Research Tissue Bank (REC Ref 11/AL/0174) for HPV related research on archive samples. Samples are available for the present project though application to the Archive Steering Committee (HPV Archive Application Ref 0034). RNA and protein were extracted from the samples using Trizol as described by the manufacturer (ThermoFischer Scientific, USA) and analysed as described.

### Tissue Microarray and Immunohistochemistry

A cervical cancer tissue microarray (TMA) containing 39 cases of cervical cancer and 9 cases of normal cervical tissue (in duplicate) were purchased from GeneTex, Inc. (GTX21468). Slides were deparaffinised in xylene, rehydrated in a graded series of ethanol solutions and subjected to antigen retrieval in citric acid. Slides were blocked in normal serum and incubated in primary antibody (USP13 (D4P3M; 12577, Cell signalling Technologies (CST)) or Mcl-1 (S-19; sc-819, Santa Cruz Biotechnology (SCBT)) overnight at 4 °C. Slides were then processed using the VECTASTAIN® Universal Quick HRP Kit (PK7800; Vector Laboratories) as per the manufacturer’s instructions. Immunostaining was visualised using 3,3’-diaminobenzidine (Vector® DAB (SK-4100; Vector Laboratories)). USP13 and Mcl-1 immunostaining quantification was automated using ImageJ with the IHC Profiler plug-in (70). Histology scores (H-score) were calculated based on the percentage of positively stained tumour cells and the staining intensity grade (71). The staining intensities were classified into the following four categories: 0, no staining; 1, low positive staining; 2, positive staining; 3, strong positive staining. H score was calculated by the following formula: (3 x percentage of strong positive tissue) + (2 x percentage of positive tissue) + (percentage of low positive tissue), giving a range of 0 to 300.

### Cell culture

HeLa (HPV18+ cervical adenocarcinoma cells), SW756 (HPV18+ cervical squamous carcinoma cells), SiHa (HPV16+ cervical squamous carcinoma cells), CaSKi (HPV16+ cervical squamous carcinoma cells) and C33A (HPV-cervical squamous carcinoma) cells obtained from the ATCC and grown in DMEM supplemented with 10% FBS (ThermoFischer Scientific, USA) and 50 U/mL penicillin. Primary normal human keratinocytes (NHKs) were maintained in serum free medium (SFM; GIBCO, UK) supplemented with 25 μg/mL bovine pituitary extract (GIBCO, UK) and 0.2 ng/mL recombinant EGF (GIBCO, UK). All cells were cultured at 37°C and 5% CO2. Cells were negative for Mycoplasma during this investigation. Cell identify was confirmed by STR profiling.

### Plasmids, siRNA and inhibitors

pRK5-FLAG-USP13 (Addgene plasmid #61741) and pcDNA3.1-V5-hMcl-1 (Addgene plasmid #25375) were purchased from Addgene (Cambridge, MA, USA). The USP13 C345A mutant was created using the Q5 Site-Directed Mutagenesis Kit (New England BioLabs Inc, MA, USA) using the following primers: F – 5’ TGGGCAACAGCGCCTATCTCAGCTC ‘3 and R – 5’ GGTTCTTCAGACCCGTGT ‘3. HA-Ubiquitin, HA-Ubiquitin (K48R) and HA-Ubiquitin (K63R) were a kind gift from Prof Paul Lehner (University of Cambridge, UK). siRNA targeting USP13 were purchased from Qiagen (FlexiTube GeneSolution GS8975 for JUN; SI03074498, SI03061968, SI00058100, SI00058093). The small molecule inhibitors MG132, JNK-IN-8, Spautin-1 and ABT-263 were purchased from Calbiochem and used at the indicated concentrations.

### Transfections and mammalian cell lysis

Transfection of plasmid DNA or siRNA was performed with a DNA to Lipofectamine® 2000 (ThermoFisher) ratio of 1:2.5. 48 or 72 hr post transfection, cells were lysed in lysis buffer for western blot analysis, or reseeded into new plates for growth curve analysis, colony formation assays or soft agar assays.

### Deubiquitination assays

FLAG-USP13, pcDNA3.1-V5-hMcl-1 and/or HA-ubiquitin plasmids were transfected into HEK293 cells using Lipofectamine 2000. After 40 h, the cells were treated with 20 μM MG132 (Calbiochem) for 8 h. The cells were then washed with PBS and lysed in HEPES buffer supplemented 100 μM N-ethylmaleimide and protease inhibitor cocktail (Roche). The lysates were centrifuged and incubated with 2 μg Mcl-1 antibody (S-19; sc-819, SCBT) at 4°C overnight with continuous rotation. A/G agarose beads were then added, and the lysate/antibody/bead mix was incubated at 4°C overnight with continuous rotation Beads were then washed in HEPES buffer and boiled in Laemmli loading buffer prior to SDS PAGE and western blot analysis with an anti-HA antibody (HA-7; H9658, Sigma).

### Immunoprecipitations

Assays were performed as previously described (72). Briefly, cells were transfected with FLAG-USP13, pcDNA3.1-V5-hMcl-1 or both. Cell lysates were harvested and then incubated with USP13, FLAG or Mcl-1 antibodies and incubated at 4°C overnight with continuous rotation. A/G agarose beads were then added, and the lysate/antibody/bead mix was incubated at 4°C overnight with continuous rotation. Beads were then washed in lysis buffer and boiled in Laemmli loading buffer prior to SDS PAGE and western blot analysis.

### Western blot analysis

Equal amounts of protein from cell lysates were separated by SDS PAGE and transferred onto a nitrocellulose membrane by a semi-dry transfer method (Trans Blot® SD Semi-Dry Transfer cell, Bio-Rad, USA). Membranes were blocked with 5 % milk solution before incubation with primary antibodies at 1:1000 dilution unless otherwise stated: USP13 (D4P3M; 12577, CST), Mcl-1 (S-19; sc-819, SCBT), Phospho-c-Jun (Ser73) (D47G9; 3270, CST), c-Jun (60A8; 9165, CST), FLAG (F1804, Sigma-Aldrich), (HA-7; H9658, Sigma) and GAPDH (SCBT; sc365062) (1:5000) as a loading control. Horseradish peroxidase (HRP)-conjugated secondary antibodies (Sigma Aldrich, USA) were used at a 1:5000 dilution. Proteins were detected using WesternBright ECL (Advansta, USA) and visualised on X-ray film.

### Chromatin immunoprecipitation

Putative AP-1 binding sites in the USP13 3’UTR were identified using the AceView program. HeLa cells were treated with JNK-IN-8 for the required incubation time. Cells were processed for ChIP analysis as previously described (73). Briefly, cells were fixed in 1% formaldehyde for 10 min at room temperature, quenched in 0.25 M glycine, and washed in ice-cold PBS. Cells were harvested by scraping and then lysed in cell lysis buffer (10 mM Tris-HCl, pH 8.0, 10 mM NaCl, 0.2% NP-40, 10 mM sodium butyrate, 50 μg/ml phenylmethylsulfonyl fluoride (PMSF), 1× complete protease inhibitor). Nuclei were collected by centrifugation at 2,500 rpm at 4 °C and resuspended in nuclear lysis buffer (50 mM Tris-HCl, pH 8.1, 10 mM EDTA, 1% SDS, 10 mM sodium butyrate, 50 μg/ml PMSF, 1× complete protease inhibitor). Extracted chromatin was then sonicated and chromatin concentration was determined. Approximately 100 μg of chromatin from each sample was used for the experiment. c-Jun was immunoprecipitated using a ChIP grade anti-c-Jun antibody (60A8; 9165, CST). A/G magnetic beads were used to pull down the antibody-chromatin complex. To show antibody specificity, each of the samples were pulled down with an IgG isotype control. The immunoprecipitated chromatin was then processed for quantitative PCR (qPCR); the primer sequences used are available on request. Fold-enrichment compared to negative control IgG isotype control was calculated as in (74).

### RNA extraction, cDNA synthesis and quantitative Real Time-PCR

RNA extraction for qRT-PCR was performed using an E.Z.N.A Total RNA Kit I (Omega bio-tek, USA). cDNA was synthesised with 1 μg of input RNA and iScript cDNA synthesis kit (Bio Rad, USA). qRT-PCR was performed on the synthesised cDNA on a Corbett Rotor-Gene 6000 using QuantiFast SYBR Green PCR kit (Qiagen, USA) and analysed using the ∆∆CT method (75) normalised to the U6 housekeeping gene. Primer sequences are available on request.

### Cell viability (MTT) assays

Cells were seeded in triplicate in 96-well plates and subjected to the indicated treatments for 48◻h before adding MTT reagent and measuring the absorbance at 57025FBnm according to the manufacturer’s instructions (Sigma, USA). For combination matrices, cells were seeded in 96-well plates. After 24◻h, cells were treated with ABT-263 (dose range of 0–525FBμM) and Spautin-1 (dose range of 0–20◻μM) in a 6◻×◻6 matrix. Cells were cultured with inhibitors for 4825FBh and cell viability was determined using MTT assay as above. The Bliss synergy heatmap and synergy score was calculated using SynergyFinder (76).

### Colony formation assay

48 or 72 hr post-treatment or transfection, cells were trypsinised and reseeded in a six well plate at 500 cells per well and left to incubate for 14 - 21 days. Colonies were then stained (1 % crystal violet, 25 % methanol) and colonies were counted manually. Each experiment was repeated a minimum of 3 times.

### Soft agar assay

Cells were treated or transfected as required. 60 mm dishes were coated with a layer of 1 % agarose (ThermoFisher Scientific) in 2X DMEM (ThermoFisher Scientific) supplemented with 20 % FBS. 48 or 72 hr post-transfection, cells were trypsinised and resuspended in 0.7 % agarose in 2X DMEM (ThermoFisher Scientific) supplemented with 20 % FBS at 1000 cells/mL. Once set, DMEM supplemented with 10 % FBS and 50 U/mL penicillin was added. The plates were then incubated for 14 - 21 days. Each experiment was repeated at least three times. Visible colonies were counted manually.

### Flow cytometry

Cells were transfected as required. 72 hr post-transfection, cells were harvested and fixed in 70 % ethanol overnight. The ethanol was removed, and cells washed with PBS containing 0.5 % (w/v) BSA. Cells were stained with PBS containing 0.5 % BSA, 50 μg/mL propidium iodide (Sigma-Aldrich) and 5 μg/mL RNase (Sigma-Aldrich) and incubated for 30 min at room temperature. Samples were processed on a CytoFLEX S flow cytometer (Beckman Coulter) and analysed using CytExpert (Beckman Coulter).

### Annexin V assay

Annexin V apoptosis assay (TACS Annexin V kit; 4830-250-K) was performed as indicated on the product datasheet. Briefly, cells seeded in 6 well plates were treated or transfected as required. Cells were trypsinised and collected by centrifugation. 1×10^6^ cells were then incubated in 100 μL Annexin V reagent (10 μL 10 x binding buffer, 10 μL propidium iodide, 1 μL Annexin V-FITC (diluted 1 in 500) and 880 μL ddH2O) for 15 mins at room temperature in the dark. Samples were diluted in 1 x binding buffer before analysis by flow cytometry. Samples were processed on a CytoFLEX S flow cytometer (Beckman Coulter) and analysed using CytExpert (Beckman Coulter).

### Statistical Analysis

Unless otherwise indicated, data was analysed using a two-tailed, unpaired Student’s t-test and graphs were prepared using the GraphPad Prism software (GraphPad, USA).

## Acknowledgements

We are particularly grateful to Prof. Roger Davis (UMASS, United States) and Yihong Ye (NIDDK, NIH, USA) for generous provision of reagents through the Addgene repository. We thank Prof. Paul Lehner (University of Cambridge) for the ubiquitin expression plasmids and Prof. Adrian Whitehouse (University of Leeds) for providing the V5 antibody. We additionally thank the Scottish HPV Investigators Network (SHINE) for providing HPV positive cytology samples. Finally, we thank Dr Stephen Griffin for critical reading of the manuscript.

---

**Supplementary Figure S1.**
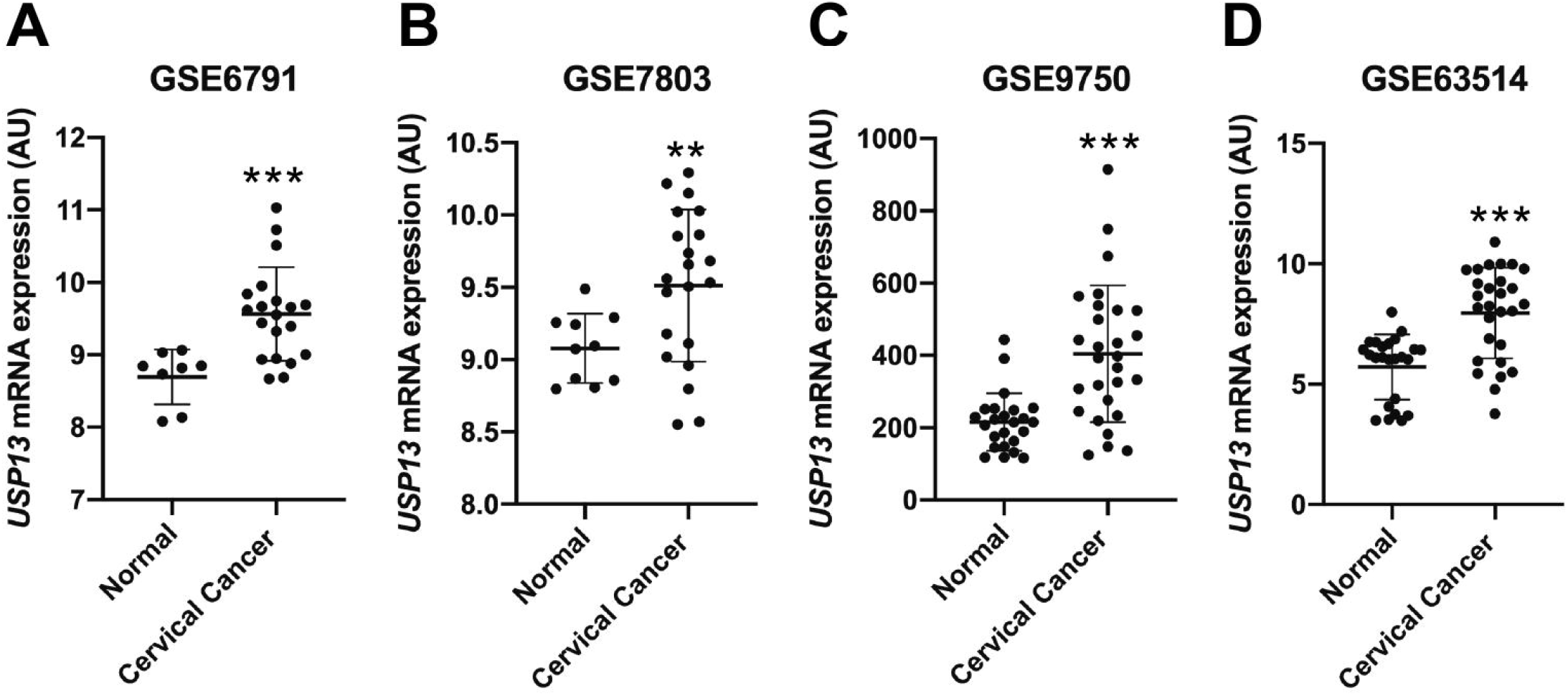
*USP13* mRNA is upregulated in cervical cancer. **A)***USP13* mRNA expression from the GEO databases GSE6791, GSE7803, GSE9750 and GSE63514. Error bars represent the mean ± standard deviation. * p<0.5; ** p<0.01; *** p<0.001 (Student’s t-test).

**Supplementary Figure S2.**
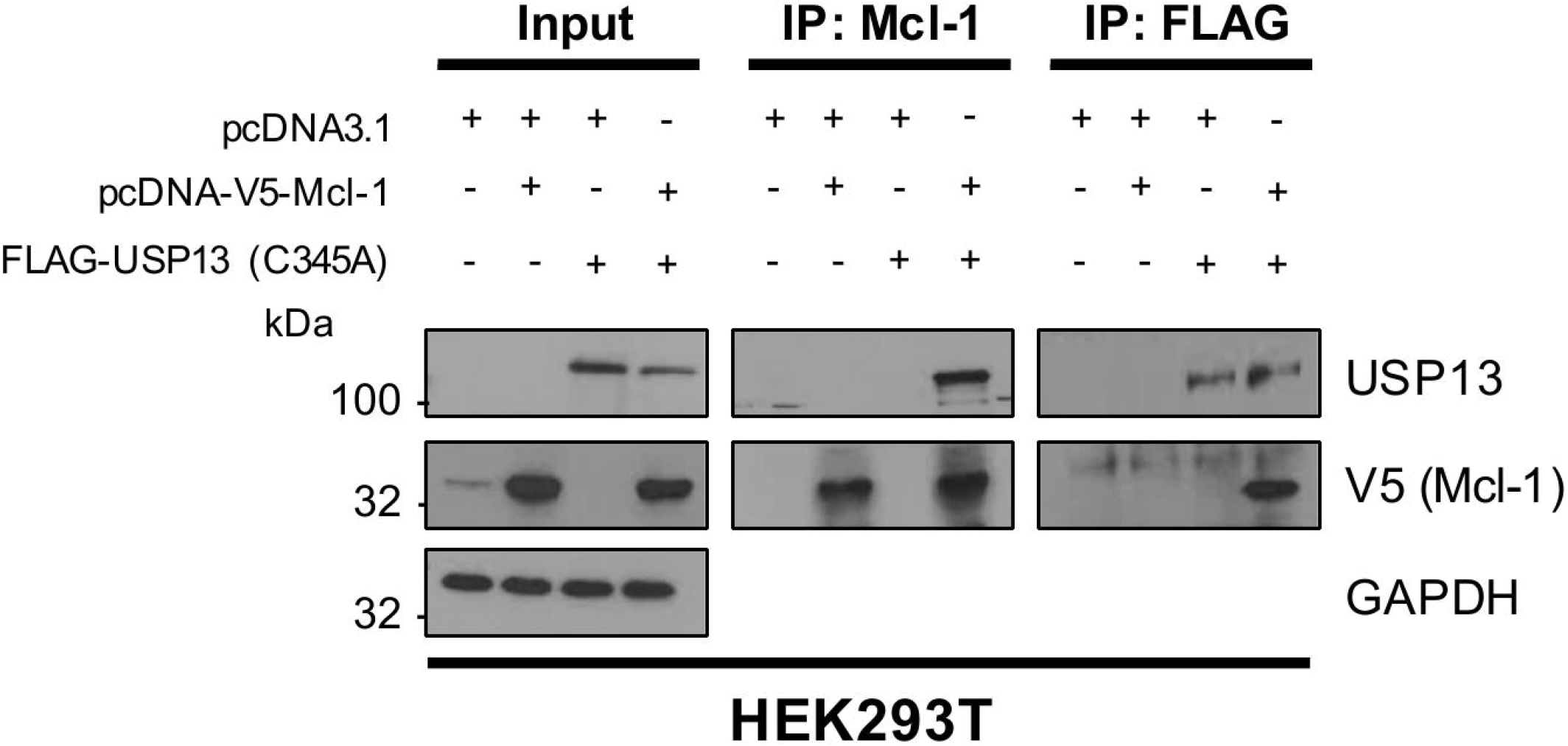
Catalytically inactive USP13 interacts with Mcl-1. **A)** HEK293T cells were transfected with Mcl-1, Flag-USP13 (C345A), or both Mcl-1 and FLAG-USP13 (C345A). Cells were treated with 10 μM MG132 for 6 hours and either Mcl-1 or USP13 were immunoprecipitated using an anti-Mcl1 or anti-FLAG antibody. Co-immunoprecipitated Mcl-1 or FLAG-USP13 were detected using the respective antibodies. GAPDH was used as a loading control.

